# A nonsense mutation of bone morphogenetic protein-15 (BMP15) causes both infertility and increased litter size in pigs

**DOI:** 10.1101/2020.08.03.222521

**Authors:** Gabriele Flossmann, Christine Wurmser, Hubert Pausch, Amabel Tenghe, Jörg Dodenhoff, Günther Dahinten, Kay-Uwe Götz, Ingolf Russ, Ruedi Fries

## Abstract

**Background:** Atypical external genitalia are often a sign of reproductive organ pathologies and infertility with both environmental or genetic causes, including karyotypic abnormalities. Genome-wide association studies (GWAS) provide a means for identifying chromosomal regions harboring deleterious DNA-variants causing such phenotypes. We performed a GWAS to unravel the causes of incidental cases of atypically small vulvae in German Landrace gilts.

**Results:** A case-control GWAS involving Illumina porcine SNP60 BeadChip-called genotypes of 17 gilts with atypically small vulvae and 1,818 control animals (fertile German Landrace sows) identified a significantly associated region on the X-chromosome (P = 8.81 × 10^-43^). Inspection of whole-genome sequencing data in the critical area allowed us to pinpoint a likely causal variant in the form of a nonsense mutation of bone morphogenetic protein-15 (Sscrofa11.1_X:g.44618787C>T, BMP15:p.R212X). The mutant allele occurs at a frequency of 6.2% in the German Landrace breeding population. Homozygous gilts exhibit underdeveloped, most likely not functional ovaries and are not fertile. Male carriers do not seem to manifest defects. Heterozygous sows produce 0.41±0.02 (P=4.5 × 10^-83^) piglets more than wildtype animals. However, the mutant allele’s positive effect on litter size accompanies a negative impact on lean meat growth.

**Conclusion:** Our results provide an example for the power of GWAS in identifying the genetic causes of a fuzzy phenotype and add to the list of natural deleterious BMP15 mutations that affect fertility in a dosage-dependent manner, the first time in a poly-ovulatory species. We advise eradicating the mutant allele from the German Landrace breeding population since the adverse effects on the lean meat growth outweigh the larger litter size in heterozygous sows.

## Background

A number of German Landrace sows with atypically small vulvae have been recently observed on several Bavarian farms. Most of the sows turned out to be anoestrous or, when they showed signs of oestrus, insemination was not possible due to problems in introducing the artificial insemination device. Usually, more than one gilt of a litter was affected by the abnormality but not all. Although environmental factors such as mycotoxin intoxication or nutritional deficiencies could not be ruled out to cause the observed condition, we presumed a genetic etiology. Abnormal external genitalia are often seen in the context of intersexuality. Karyotypically female pigs (38, XX) have been reported with phenotypes ranging from complete external masculinization with testes to an external female appearance with enlarged clitorises, testes, and ovotestes [1]. A genome-wide association study pointed to the causal implication of *SOX9* in the sex-reversal phenotype of these SRY-negative XX pigs [2]. To explore possible genetic causes of the newly observed phenotype of abnormal female genitalia we also undertook a genome-wide case-control study that led us to a nonsense mutation in *BMP15*, most likely the cause for the abnormality in homozygously affected animals but also for increased litter size in heterozygous sows.

## Results

### Genome-wide association study

We had access to tissue samples of 43 German Landrace gilts, considered as affected based on their abnormally small vulvae. The 43 affected animals originated from 21 litters of 20 sows and 11 known boars and were from four breeding units; the sire of one litter was unknown (Additional file 1). Samples from 17 affected individuals were genotyped on the Illumina Porcine SNP60 BeadChip and subjected as cases to a genome-wide association study (GWAS) with 1,818 fertile sows from the Bavarian German Landrace population as controls. Association testing revealed a cluster of 18 significantly associated SNPs (P < 9.87 × 10^-7^) between 38.6 Mb and 108.7 Mb on the X-chromosome (Figure 1a). The most significant SNP (P = 8.81 × 10^-43^) on the X-chromosome (rs328794518, 88,499,366 bp, Sscrofa11.1 assembly) was an intron variant in the *ENSSSCG00000012566* gene. We considered significantly associated, scattered SNPs on other chromosomes as artifacts due to the inflation of significant signals (λ = 1.26, Figure 1b).

**Figure 1:**
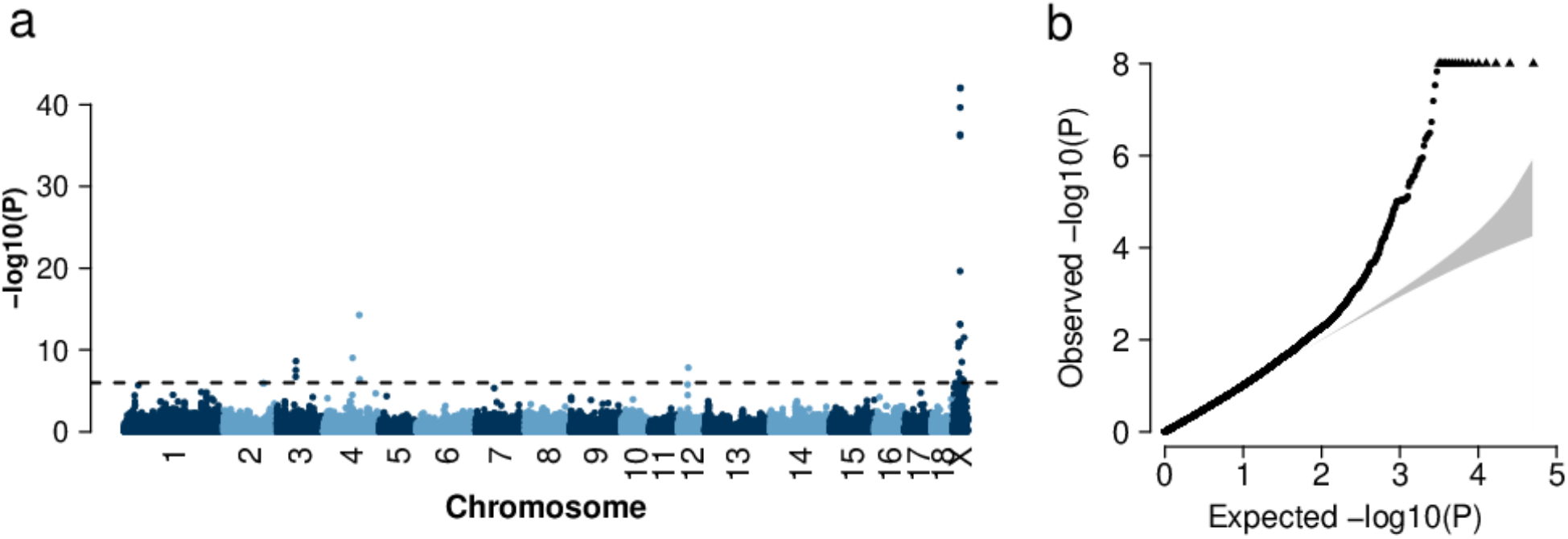
Genome-wide association study for atypically small vulvae in 1835 German Landrace sows. Manhattan plot (**a**) representing the association of 50,649 SNPs with atypically small vulvae in the German Landrace population. The dots above the dashed line represent SNPs with P-values less than 9.87 × 10^-7^ (Bonferroni corrected significance threshold) and Quantile-quantile plot (**b**). The grey shaded area represents the 95% concentration band under the null hypothesis of no association. Triangles represent SNPs with P-values less than 9.87 × 10^-7^.

### Identification of the underlying mutation by analyzing whole-genome re-sequencing (WGS) data

Re-sequencing data of 42 pigs (22 German Landrace and 20 Piétrain boars and fertile sows) were available for identifying possible causal variants. The sire of eight affected gilts was among the 42 sequenced pigs. We located 828 coding variants within the associated region (38.6 Mb to 108.7 Mb), of which 366 were missense mutations, 30 were frameshift variants, and five were nonsense-mutations. The sire of the affected gilts was hemizygous for one of the five nonsense-mutations. This nonsense-mutation, with no previous reports, is a C to T substitution in the second exon of *BMP15* (Sscrofa11.1_X:g.44618787C>T, NP_001005155.2:p.R212X). In addition to the sire of affected gilts, two German Landrace boars and one German Landrace sow were carriers of the nonsense-mutation (Additional file 2). None of the Piétrain pigs carried the mutation.

### Validation of the nonsense mutation in *BMP15*

All 22 German Landrace pigs with whole-genome sequences were genotyped by Sanger sequencing to confirm the mutation technically (Additional file 3). Complete concordance with WGS derived genotypes corroborates the technical validity of the variant. Next, we genotyped 43 sows, assessed by the farmers to exhibit small vulvae, 29 full siblings with normal vulvae, and 4,869 fertile sows, using either a KASP™ genotyping assay or a new customized version of the Illumina Porcine SNP60 BeadChip containing the p.R212X variant of BMP15. Of the 43 sows with abnormal vulvae, 36 were homozygous for the T-allele, three were heterozygous (C/T), and four were homozygous for the wild type allele (C). Of the seventeen animals that we considered as cases in the GWAS, only eleven turned out to be homozygous for the mutant allele; three animals where heterozygous and three animals were homozygous for the wildtype allele. None of the 4,869 fertile sows and none of the 29 unaffected full siblings of affected gilts were homozygous for the T-allele (Table 1). The frequency of the T-allele in the 4,869 fertile sows was 0.062.

**Table 1:** Genotypes of the p.R212X variant of BMP15 for 4,941 German Landrace sows

| Genotype | Sows with small vulvae | Full siblings with normal vulvae | Fertile German Landrace sows |
| --- | --- | --- | --- |
| **C/C** (ref/ref) | 4 | 0 | 4,263 |
| **C/T** (ref/alt) | 3 | 29 | 606 |
| **T/T** (alt/alt) | 36 | 0 | 0 |
| **N sows** | 43 | 29 | 4,869 |

While fertile sows and gilts that were classified to have normal-sized vulvae were not homozygous for the nonsense mutation of *BMP15*, seven of the 43 presumably affected gilts carried one or two wild type alleles. This finding is not compatible with the hypothesis of the nonsense mutation causing small vulvae. However, assessing the size of the vulva is subjective, and the discrimination between affected and unaffected animals is prone to error. On the other hand, if an animal’s status is assessed based on the objective criterium of fertility (i.e., an animal having one or more litters), the hypothesis that the nonsense mutation of *BMP15* causes underdeveloped external sexual organs and consequent infertility remains valid.

### Effect of the mutation on reproductive organs

The initial phenotyping consisted of on-farm observations of gilts with underdeveloped vulvae. A more thorough assessment of the phenotype in the field turned out to be impracticable. Therefore, we acquired two gilts that were heterozygous for BMP15:p.R212X and mated them on an experimental farm with a boar carrying the mutation. One of the resulting litters consisted of male piglets only, while the other litter consisted of thirteen piglets, ten of them female. Of the female piglets, five were homozygous for the T-allele, and five were heterozygous (C/T). At the age of six months, eight of the ten female pigs (five homozygous and three heterozygous) were slaughtered, and their uteri and ovaries resected. The uterus horns of the homozygous animals were considerably smaller, specifically less voluminous (Figure 2a, b, c) or less coiled (Figure 2d and e) than those of their heterozygous full siblings (Figure 2f, g, h). The differences in the appearance of the ovaries were even more pronounced. As shown in Figure 3, the ovaries of homozygous gilts (Figure 3a to e) did not exhibit follicles, whereas multiple follicles were evident in the ovaries of their heterozygous full siblings (Figure 3f to h). Besides the morphologically assessed underdevelopment of the ovaries and uteri, there may also be functional deficits as indicated by cysts in two homozygously affected gilts (Figure 3d and e). It is noteworthy that the two gilts with cysts exhibit uterus horns that are more voluminous (Figure 2d and e) than those of their homozygous siblings (Figure 2a, b, c), but less coiled than those of their heterozygous siblings (Figure 2f to h).

**Figure 2:**
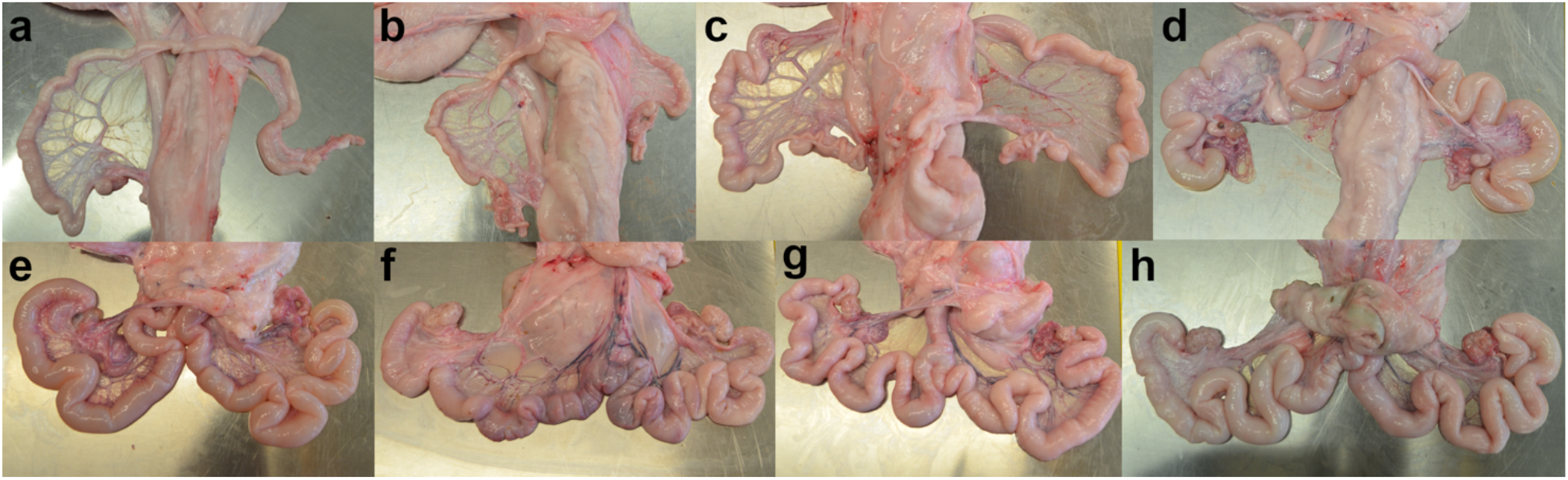
Uteri of 8 fullsibs at the age of 6 months. Uteri of fullsib gilts from the mating of a sow with genotype C/T of Sscrofa11.1_X:g.44618787C>T (BMP15:R212X) and a boar carrying the T-allele. Pictures **a** to **e** show uteri of gilts with genotype T/T and pictures **f** to **h** are from sows with genotype C/T. Note that the uterus horns shown in **d** and **e** are more voluminous than in **a**, **b** and **c**, but less coiled than those in **e**, **f** and **g**.

**Figure 3:**
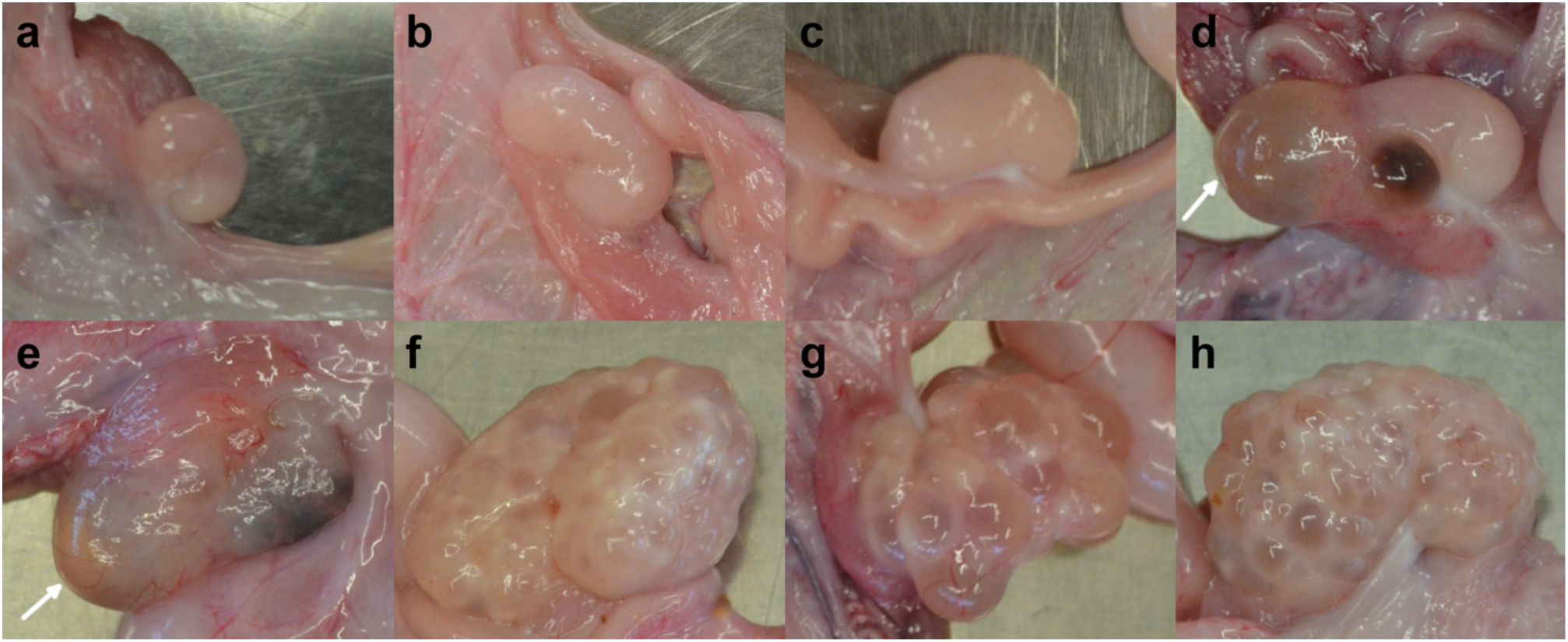
Ovaries of 8 full sibs at the age of 6 months. Ovaries of fullsib gilts from the mating of a sow with genotype C/T Sscrofa11.1_X:g.44618787C>T (BMP15:R212X) and a boar carrying the T-allele. Pictures **a** to **e** show ovaries of gilts with genotype T/T and pictures **f** to **h** are from sows with genotype C/T. The arrows point to cysts.

### Effects of the mutation on fertility and other economically important breeding traits

Nonsense and missense mutations of BMP15 increase ovulation rate in heterozygous ewes but cause infertility in ewes that are homozygous for the mutant allele (first reported by Galloway et al. (2000) [3]). Therefore, we analyzed the effects of the *BMP15* nonsense mutation (i. e., the T-allele) on fertility, measured as the number of live-born piglets, and several other economically important traits, most of them growth-related. Since genotypes were not available for all sows with breeding values, we derived the T-allele dosage based on the parents’ genotypes and the T-allele frequencies in male and female animals. A total of 5,263 genotyped parents were available (4,263 C/C-sows, 606 C/T-sows, 367 C-boars, and 27 T-boars; T/T-females cannot be parents as they are infertile). The frequency of the T-allele amounts to q_m_ = 0.069 in male, and q_f_ = 0.062 in female parents. Table 2 lists the expected dosages and Table 3 the number of animals in each dosage group.

**Table 2:** Dosage of the T-allele (causing the nonsense mutation) based on the genotypes of the parents

|  | C/C-sow | C/T-sow | C/N-sow |
| --- | --- | --- | --- |
| **C-boar** | 0 | 0.5 | $q_f = 0.062$ |
| **T-boar** | 1 | 1 | 1 |
| **N-boar** | $q_m = 0.069$ | $q_m + 0.5 = 0.569$ | $q_m + q_f = 0.131$ |

$q_m$ : T-allele frequency in boars; $q_f$ : T-allele frequency in sows; C: wildtype allele; T: mutant allele; N: allele not known.

**Table 3:** Number of sows in T-allele dosage groups

|  | T-allele dosage |  |  |  |  |  |  |
| --- | --- | --- | --- | --- | --- | --- | --- |
|  | 0.000 | 0.062 | 0.069 | 0.131 | 0.500 | 0.569 | 1.000 |
| **N sows** | 8,400<br>(4,156) | 9,623<br>(9,623) | 1,092<br>(1,092) | 9,969<br>(9,969) | 1,185<br>(1,185) | 565<br>(565) | 1,185<br>(579) |

Numbers in parentheses indicate animals with allele dosage derived from the parents' genotypes.

We applied a linear regression model to estimate the effect of the T-allele of the *BMP15* nonsense mutation on breeding values (Table 4). The T-allele has a significantly positive effect on the number of live-born piglets (+0.41 piglets). Thus, the *BMP15* mutation increases the litter size, probably as a result of an increased ovulation rate, and causes infertility in a dosage-sensitive manner (one T-allele increases the litter size, two T-alleles cause infertility). The positive effect of the *BMP15* mutation on litter size is contrasted by the significantly negative effect on traits of lean meat growth (lean meat content, belly meat content, loin eye area, loin to fat ratio) and the positively correlated feed conversion ratio. Intramuscular fat content is negatively correlated with lean meat growth and thus positively affected by the *BMP15* mutation.

**Table 4:** Estimated effects of the p.R212X-mutation on economically important traits

| Trait | N sows | Effect<br>± std. error | P-value | Signifi-<br>cance |
| --- | --- | --- | --- | --- |
| **Live-born piglets (N)** | 20,549 | +0.411 ± 0.021 | $4.50 \times 10^{-83}$ | **** |
| **Feed conversion ratio (kg/kg)** | 10,072 | -0.028 ± 0.004 | $7.60 \times 10^{-15}$ | **** |
| **Daily gain (g/day)** | 10,072 | -0.847 ± 1.684 | $6.15 \times 10^{-01}$ | |
| **Lean meat content (%)** | 10,072 | -0.889 ± 0.059 | $2.50 \times 10^{-51}$ | **** |
| **Belly meat content (%)** | 10,072 | -0.906 ± 0.057 | $2.21 \times 10^{-56}$ | **** |
| **Loin eye area (cm<sup>2</sup>)** | 10,072 | -1.216 ± 0.097 | $1.12 \times 10^{-35}$ | **** |
| **Loin to fat ratio (cm<sup>2</sup>/ cm<sup>2</sup>)** | 10,072 | -0.031 ± 0.002 | $1.06 \times 10^{-51}$ | **** |
| **Carcass length (cm)** | 10,072 | -0.030 ± 0.045 | 5.13 × 10<sup>-01</sup> |  |
| **Intramuscular fat content (%)** | 10,072 | 0.068 ± 0.007 | 6.20 × 10<sup>-25</sup> | **** |
| **pH1** | 10,072 | 0.002 ± 0.002 | 1.16 × 10<sup>-01</sup> |  |
| **Drip loss (%)** | 10,072 | -0.005 ± 0.004 | 2.16 × 10<sup>-01</sup> |  |

N sows: number of sows with a breeding value reliability ≥ 40%; Bonferroni corrected significance for eleven tests is P-value ≤ 4.5 × 10<sup>-03</sup>. \*\*\*\* significant at P ≤ 1 × 10<sup>-04</sup>

## Discussion

Identifying genetic determinants of subjectively assessed phenotypes, such as gilts’ vulvae size, seems hardly feasible. However, comparing the genotypes of seventeen gilts exhibiting relatively small vulvae with those of fertile sows enabled us to identify a distinct region on the X-chromosome to carry a potential causal mutation in a genome-wide association study. The availability of whole-genome sequence information of ancestors of the affected animals enabled us to readily identifying a likely causal variant, i.e. a nonsense mutation in *BMP15*.

Only eleven of the seventeen animals with apparently small vulvae that we considered as cases in the GWAS turned out to be homozygous for the mutant allele. This finding demonstrates the power of GWAS for identifying genetic causes even of a poorly phenotype, such as vulva size. Several factors affect their appearance. Vulvae are smaller in pre-than in postpubescent gilts, and the age of puberty varies considerably between animals [4]. A more reliable assessment of the vulva size as a possible indicator of follicular development would require the inspection of postpubescent gilts [5], which was not feasible for the present study.

BMP15 has fundamental roles in ovarian function. Its actions include (1) promotion of follicle growth and maturation; (2) regulation of the sensitivity of granulosa cells (GC) to follicle stimulation hormone (FSH); (3) prevention of GC apoptosis; (4) promotion of oocyte developmental competence and (5) determination of ovulation quota [6]. Natural mutations of *BMP15* cause primary ovarian insufficiency (POI) in women (reviewed by Patiño et al., 2017) [7]). Natural missense and nonsense mutations of ovine *BMP15* cause both an increased ovulation rate and infertility, depending on the dosage of the mutant allele [3, 8–12]. Notable exceptions are missense mutations of *BMP15* in the Grivette and Olkuska sheep breeds, as they do not cause infertility. They increase the ovulation rate and the litter size in a dosage-dependent manner without causing infertility [12]. BMP15 acts through homodimerization of the mature protein and heterodimerization with GDF9 (growth differentiation factor 9), another member of the transforming growth factor β (TGF-β) family [13]. Mutations of *GDF9* also affect fertility in sheep in the same mode as mutations of *BMP15* [8, 14, 15]. A mutation of the bone morphogenetic protein IB receptor *(BMPR1B)*, finally, causes the high prolificacy Booroola phenotype [16–18] without a negative effect in the homozygous state.

While mutations of *BMP15* critically affect follicular development and ovulation rate in dosage-sensitive manners in the mono-ovulatory human and ovine species, the effects of such mutations seem to be less pronounced in the poly-ovulatory species mouse. *BMP15*-knockout female mice have minimal ovarian histopathological defects but are subfertile due to a decreased ovulation rate [19]. However, knockdown of *BMP15* in the poly-ovulatory species swine completely inhibits ovarian follicular development, leading to infertility [20]. This latter study did not allow to assess the effect of the null allele’s heterozygous state on the ovulation rate and litter size. The natural nonsense mutation of *BMP15* that we describe in the present study is most likely a null allele as it results in a severely truncated protein (47.5% are missing) and / or nonsense-mediated mRNA decay. Our data indicate that heterozygosity of a *BMP15* mutation also enhances litter size in poly-ovulatory swine, just as has been amply reported in mono-ovulatory sheep. *GDF9* and *BMP15* mutations also seem to be more frequent in mothers of dizygotic twins than mothers of single infants or monozygotic twins [21], pointing to the possibility of an increased ovulation rate due to heterozygosity of deleterious variants in the human species as well.

Galloway et al. (2000) [3] set out to explain the paradoxical dosage effect of BMP15 and GDF9 mutations. They hypothesized that 50% of normal levels of the proteins might reduce granulosa cell mitosis and delay suppressive effects on plasma follicle-stimulating hormone (FSH) concentrations and that the concomitant reduction of the amount of steroid or inhibin would lead to the ovulation of an additional oocyte in mono-ovulatory species (or additional oocytes in poly-ovulatory species). McNatty et al. (2009) [22] showed that an earlier acquisition of responsiveness to the luteinizing hormone (LH) by granulosa cells in a higher proportion of follicles accounts for the higher ovulation-rate in heterozygous carriers of a BMP15 mutation.

To increase the litter size is an objective of pig breeding. Breeding gilts that are heterozygous for the nonsense mutation of the *BMP15* mutation could expedite the breeding for increased litter size. In detail, German Landrace boars carrying the mutation could be mated with German Large White sows. Homozygous female littermates of these boars can be diagnosed by a DNA-test and directed to fattening. All crossbred gilts sired by these boars would be carriers of the mutation. Symptomatic homozygous crossbred progeny would not result in this crossbreeding scheme since German Large White animals do not carry the *BMP15* mutation (data not shown). When mated with Piétrain boars, crossbred sows will produce an increased number of piglets suitable for fattening.

The outlined selection scheme to achieve an increased litter size is only advisable if the *BMP15* mutation does not negatively affect other economically relevant traits. However, our data reveal a significantly negative effect of the *BMP15* mutation on traits of lean meat growth. We did not expect such an effect since the *BMP15* gene is expressed exclusively in oocytes, and the protein does not seem to have any other function than to control the development of follicles and ovulation [7, 21]. However, an indirect effect of a linked variant on traits of lean meat growth cannot formally be excluded. When taking the economic weights [23] for significantly affected traits into account, we can assess the effect of the *BMP15* mutation economically. Each additional live-born piglet has a monetary value of 3.98 €, each kg less feed consumed per kg gain 22.70 €, and each percentage point of lean meat content 1.66 €, yielding a total monetary effect of the mutant allele of −0.48 € per finisher pig. Thus, using the *BMP15* mutation to increase litter size is not advisable, and the eradication of the mutant allele from the German Landrace population, if carefully executed with regard to the genetic diversity, should not cause losses.

## Conclusions

The observation of gilts with atypically small vulvae prompted a genome-wide association study that led to the detection of an associated region on the X-chromosome. The inspection of whole-genome sequencing data allowed us to pinpoint a nonsense mutation of *BMP15* (Sscrofa11.1_X:g.44618787C>T, NP_001005155.2:p.R212X) as the most likely causal variant. Like mutations of the ovine *BMP15*, the porcine mutation affects female fertility in a dosage-dependent manner. Homozygous female carriers of the variant lack proper ovary function and are infertile; heterozygous sows produce +0.41 piglets per litter. Male animals are not afflicted. Thus, our findings show that the *BMP15* nonsense mutation has similar effects in the poly-ovulatory species swine, as observed for deleterious *BMP15* mutations in the mono-ovulatory species sheep. The negative effect of the mutant allele on lean meat growth precludes using the variant for increasing the litter size by selective breeding. The recommendation is, therefore, to eradicate the *BMP15* allele from the German Landrace breed.

## Material and Methods

### DNA extraction and genotyping

DNA was extracted from tissue samples of affected animals, collected at slaughter with the DNeasy Blood and Tissue Kit (Qiagen, Germany). Genome-wide genotypes of these samples and control animals were obtained with the Illumina Porcine SNP60 BeadChip, using default parameters of Illumina’s BeadStudio for genotype calling. The SNPs’ chromosomal positions were according to the Sscrofa11.1 assembly [24]. Sanger-sequencing was used to validate the genotypes of the Sscrofa11.1_X:g.44618787C>T candidate causal variant as derived from next-generation sequencing. Genomic PCR products obtained with primers 5’-CGCCATCAACTTCACCTAGC-3’ (forward) and 5’-TCTGGGAAGAAGTTTGGCCT-3’ (reverse) were sequenced using the BigDye^®^ Terminator v1.1 Cycle Sequencing Kit (Thermo Fisher Scientific, Massachusetts, USA) on an ABI 3130xl Genetic Analyzer (Life Technologies, Carlifornia, USA) instrument. We customized a KASP™ genotyping assay (LGC Limited, UK) for typing the variant at the population level, using 5’-TCACAAGGGGCGCAGGGTTCTA-3’ as common PCR primer, 5’-GACACATGAAGCGGAGTCG-3’ as primer for the wild type C-allele, and 5’-GCTGACACATGAAGCGGAGTCA-3’ as primer for the mutant T-allele.

### Genome-wide association study (GWAS)

We retained Porcine SNP60 BeadChip derived genotypes of SNPs positioned on the autosomes and the X-chromosome if the calling rates were higher than 90% and the minor allele frequency above 1% after quality control with PLINK v. 1.9 [25]. The final data set for the GWAS comprised 50,649 SNPs, 17 case-animals, and fertile 1,818 control-animals, i.e. sows with at least one litter. A linear mixed model assessed the association between a SNP and the phenotype (0, control; 1, case) by single-locus regression using the GCTA software [26] and the following model:

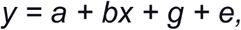

where *y* is the phenotype (coded as 1 for affected and 0 for unaffected), *a* is the mean term, *b* is the additive effect (fixed effect) of the candidate SNP to be tested for association, *x* is the SNP genotype indicator variable coded as 0, 1 or 2, *g* is the polygenic effect (random effect), i.e., the accumulated effect of all SNPs (as captured by the GRM calculated using all SNPs) and *e* is the residual. We considered SNPs with P-values less than 9.87 × 10^-7^ as significantly associated (5% Bonferroni-corrected significance threshold for 50,649 independent tests). The inflation factor was calculated using the estlambda function of the R-package GenABEL [27].

### Whole-genome sequencing

The genomes of 42 influential ancestors of two pig populations (22 German Landrace: 14 boars and eight sows, 20 Piétrains: 18 boars and two sows) were sequenced to an average coverage of 12.68x (minimum: 7.90x, maximum: 16.05x). The sire of eight sows with atypically small vulvae had a coverage of 13.95x. Library preparation followed ultrasonic fragmentation (Covaris: 50 s, 5% duty factor) using the Illumina TruSeq DNA PCR-Free Sample Preparation Kit with an insert size of 350 bp. The libraries’ sequencing was in the 125 bp paired-end mode on the HiSeq2500 instrument (Illumina, San Diego, USA). The reads were aligned to the reference sequence (Sscrofa11.1) using the Burrow-Wheeler aligner (BWA) [28]. Variants were called with GATK [29] and visualized with the Integrative Genomics Viewer (IGV) [30]. The Ensembl Variant Effect Predictor (VEP) [31] was used to annotate the variants according to the RefSeq (v. 2017_05) annotation of the Sscrofa11.1-assembly.

### Mating of animals carrying the mutation and inspection of ovaries and uteri in female offspring

Two gilts carrying the nonsense-allele of the BMP15:pR212X variant were acquired and raised on the experimental station Thalhausen of the Technical University of Munich. The gilts exhibited regular estrous cycles and were artificially inseminated at eight months’ of age, with the semen of a boar also carrying the mutation. After normal gestation, one sow gave birth to only male piglets that we did not further consider for the study, and the other sow delivered thirteen piglets, ten of them female. Weaning of the litter was at the age of four weeks. The ten females stayed in a standard pen under standard rearing conditions until they reached three months and were then divided into two groups with five pigs each and further kept under standard conditions. We genotyped the animals at Sscrofa11.1_X:g.44618787C>T with the KASP™-assay (see above) after blood samples were drawn at the age of 8 weeks and DNA was extracted by proteinase K treatment and the salting-out method. At six months of age, eight of the ten female pigs (five with T/T- and three with the C/T-genotype) were slaughtered. Uteri and ovaries were removed and examined.

### Deriving the T-allele’s dosage of Sscrofa11.1_X:g.44618787C>T in sows without genotype information

As genotypes were not available for all sows with breeding values, we derived genotypes from the parents’ genotypes to increase the number of informative animals for studying the effects of the mutation. When taking the X-chromosomal inheritance of the *BMP15* mutation (Sscrofa11.1_X:g.44618787C>T) into account, the following applies for female progeny: Parents that are C/C (sow) and C (boar) have C/C-progeny exclusively (dosage of T equals 0); if the mother is C/T and the boar is C, the progeny’s dosage of T equals 0.5. If the boar is C and the mother is not directly genotyped, i.e., either C/T or C/C (not T/T, as this genotype is not compatible with fertility), the dosage of the T-allele in the progeny amounts to the T-allele’s frequency in the female population. If the boar’s genotype is T, all resulting fertile females are C/T (dosage of T equals 1). If the boar has no genotype and the mother is C/C, the dosage of T in the progeny corresponds to the T-allele’s frequency in the male population. If the mother is C/T, the resulting dosage of T in the progeny is 0.5 plus the male T-allele’s frequency. Finally, if the boar and the sow lack direct genotype information, i.e., the sow is either C/C or C/T, the female progeny’s T-allele dosage is the sum of the male and the female T-allele’s frequency.

### Estimating the T-allele’s effect on breeding values

We used the linear regression model of the statsmodels package [32] in the Python environment (v. 3.7 [33]) to estimate the effects of the mutant T-allele on breeding values:

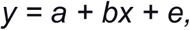

where *y* is estimated breeding value, *a* is the mean term, *b* is the additive effect of the T-allele, *x* is the T-allele dosage (0.000, 0.062, 0.069, 0.131, 0.500, 0.569, 1.00), and *e* is the residual. An effect was considered significant if P ≤ 0.0045 (the Bonferroni corrected significance level for eleven tests, i.e., breeding values). We considered only breeding values with a reliability of at least 40%.

## List of abbreviations

BMP15: Bone morphogenetic protein 15
C-boar: boar hemizygous for the wildtype allele of the BMP15:p.R212X-mutation
C/C-sow: sow homozygous for the wildtype allele of the BMP15:p.R212X-mutation
C/N-sow: sow with unknown genotype for the BMP15:p.R212X-mutation
C/T-sow: sow carrying the BMP15:p.R212X-mutation in the heterozygous state
GWAS: Genome wide association study
N-boar: boar with unknown genotype for the BMP15:p.R212X-mutation
NM_001005155.2: c.687C>T (BMP15:c687C>T)
NP_001005155.2: p.R212X (BMP15:p.R212X)
SNP: Single nucleotide polymorphism
Sscrofa11.1_X: g.44618787C>T
T-boar: boar hemizygous for the mutant allele of the BMP15:p.R212X-mutation

## Declarations

### Ethics approval and consent to participate

Ethics approval was obtained by the Regierung von Oberbayern (Sachgebiet 54) for taking blood samples from the female piglets of carrier matings and slaughter for examining the uteri and ovaries. Ethical approval was not necessary otherwise because analyses were performed on existing data obtained as part of routine data recording for the Bavarian pig breeding association.

### Consent for publication

All authors have seen the manuscript and agree with the contents.

### Availability of data and materials

All relevant data are included within the article and its additional files.

### Authors’ contributions

GF performed molecular-genetic analyses. CW and RF performed next generation sequencing and variant detection. GF, HP, AT and RF analyzed data. JD, GD and KUG provided breeding values and information on affected animals. IR provided genotype data. RF, HP and GF conceived the study. GF and RF wrote the manuscript.

## Acknowledgements

We thank the owners of the affected animals for communicating their observations.

## References

1. Pailhoux E, Pelliniemi L, Barbosa A, Parma P, Kuopio T, Cotinot C. Relevance of intersexuality to breeding and reproductive biotechnology programs; XX sex reversal in pigs. Theriogenology. 1997;47:93–102.

2. Rousseau S, Iannuccelli N, Mercat M-J, Naylies C, Thouly J-C, Servin B, et al. A genome-wide association study points out the causal implication of SOX9 in the sex-reversal phenotype in XX pigs. PLoS ONE. 2013;8:e79882.

3. Galloway SM, McNatty KP, Cambridge LM, Laitinen MP, Juengel JL, Jokiranta TS, et al. Mutations in an oocyte-derived growth factor gene (BMP15) cause increased ovulation rate and infertility in a dosage-sensitive manner. Nat Genet. 2000;25:279–83.

4. Knauer MT, Cassady JP, Newcom DW, See MT. Phenotypic and genetic correlations between gilt estrus, puberty, growth, composition, and structural conformation traits with first-litter reproductive measures. J Anim Sci. 2011;89:935–42.

5. Graves KL, Mordhorst BR, Wright EC, Hale BJ, Stalder KJ, Keating AF, et al. Identification of measures predictive of age of puberty onset in gilts. Trans Anim Sci. 2020;4:285–92.

6. Persani L, Rossetti R, Di Pasquale E, Cacciatore C, Fabre S. The fundamental role of bone morphogenetic protein 15 in ovarian function and its involvement in female fertility disorders. Hum Reprod Update. 2014;20:869–83.

7. Patiño LC, Walton KL, Mueller TD, Johnson KE, Stocker W, Richani D, et al. BMP15 Mutations Associated With Primary Ovarian Insufficiency Reduce Expression, Activity, or Synergy With GDF9. J Clin Endocrinol Metab. 2017;102:1009–19.

8. Hanrahan JP, Gregan SM, Mulsant P, Mullen M, Davis GH, Powell R, et al. Mutations in the genes for oocyte-derived growth factors GDF9 and BMP15 are associated with both increased ovulation rate and sterility in Cambridge and Belclare sheep (Ovis aries). Biol Reprod. 2004;70:900–9.

9. Bodin L, Di Pasquale E, Fabre S, Bontoux M, Monget P, Persani L, et al. A Novel Mutation in the Bone Morphogenetic Protein 15 Gene Causing Defective Protein Secretion Is Associated with Both Increased Ovulation Rate and Sterility in Lacaune Sheep. Endocrinology. 2007;148:393–400.

10. Martinez-Royo A, Jurado JJ, Smulders JP, Martí JI, Alabart JL, Roche A, et al. A deletion in the bone morphogenetic protein 15 gene causes sterility and increased prolificacy in Rasa Aragonesa sheep. Anim Genet. 2008;39:294–7.

11. Monteagudo LV, Ponz R, Tejedor MT, Laviña A, Sierra I. A 17 bp deletion in the Bone Morphogenetic Protein 15 (BMP15) gene is associated to increased prolificacy in the Rasa Aragonesa sheep breed. Anim Reprod Sci. 2009;110:139–46.

12. Demars J, Fabre S, Sarry J, Rossetti R, Gilbert H, Persani L, et al. Genome-wide association studies identify two novel BMP15 mutations responsible for an atypical hyperprolificacy phenotype in sheep. PLoS Genet. 2013;9:e1003482.

13. Shimasaki S, Moore RK, Otsuka F, Erickson GF. The Bone Morphogenetic Protein System In Mammalian Reproduction. Endocr Rev. 2004;25:72–101.

14. Nicol L, Bishop SC, Pong-Wong R, Bendixen C, Holm L-E, Rhind SM, et al. Homozygosity for a single base-pair mutation in the oocyte-specific GDF9 gene results in sterility in Thoka sheep. Reproduction. 2009;138:921–33.

15. Silva BDM, Castro EA, Souza CJH, Paiva SR, Sartori R, Franco MM, et al. A new polymorphism in the Growth and Differentiation Factor 9 (GDF9) gene is associated with increased ovulation rate and prolificacy in homozygous sheep. Anim Genet. 2011;42:89–92.

16. Mulsant P, Lecerf F, Fabre S, Schibler L, Monget P, Lanneluc I, et al. Mutation in bone morphogenetic protein receptor-IB is associated with increased ovulation rate in Booroola Mérino ewes. Proc Natl Acad Sci USA. 2001;98:5104–9.

17. Souza CJ, MacDougall C, MacDougall C, Campbell BK, McNeilly AS, Baird DT. The Booroola (FecB) phenotype is associated with a mutation in the bone morphogenetic receptor type 1 B (BMPR1B) gene. J Endocrinol. 2001;169:R1–6.

18. Wilson T, Wu XY, Juengel JL, Ross IK, Lumsden JM, Lord EA, et al. Highly prolific Booroola sheep have a mutation in the intracellular kinase domain of bone morphogenetic protein IB receptor (ALK-6) that is expressed in both oocytes and granulosa cells. Biol Reprod. 2001;64:1225–35.

19. Yan C, Wang P, DeMayo J, DeMayo FJ, Elvin JA, Carino C, et al. Synergistic roles of bone morphogenetic protein 15 and growth differentiation factor 9 in ovarian function. Mol Endocrinol. 2001;15:854–66.

20. Qin Y, Tang T, Li W, Liu Z, Yang X, Shi X, et al. Bone Morphogenetic Protein 15 Knockdown Inhibits Porcine Ovarian Follicular Development and Ovulation. Front Cell Dev Biol. 2019;7:286.

21. Belli M, Shimasaki S. Molecular Aspects and Clinical Relevance of GDF9 and BMP15 in Ovarian Function. Vitam Horm. 2018;107:317–48.

22. McNatty KP, Heath DA, Hudson NL, Lun S, Juengel JL, Moore LG. Gonadotrophin-responsiveness of granulosa cells from bone morphogenetic protein 15 heterozygous mutant sheep. Reproduction. 2009;138:545–51.

23. Zuchtziele. https://www.lfl.bayern.de/itz/schwein/046844/index.php. Accessed 26 Jun 2020.

24. Warr A, Affara N, Aken B, Beiki H, Bickhart DM, Billis K, et al. An improved pig reference genome sequence to enable pig genetics and genomics research. Gigascience. 2020;9.

25. Chang CC, Chow CC, Tellier LC, Vattikuti S, Purcell SM, Lee JJ. Second-generation PLINK: rising to the challenge of larger and richer datasets. Gigascience. 2015;4:7.

26. Yang J, Lee SH, Goddard ME, Visscher PM. GCTA: a tool for genome-wide complex trait analysis. Am J Hum Genet. 2011;88:76–82.

27. Aulchenko YS, Ripke S, Isaacs A, van Duijn CM. GenABEL: an R library for genome-wide association analysis. Bioinformatics. 2007;23:1294–6.

28. Li H, Durbin R. Fast and accurate short read alignment with Burrows-Wheeler transform. Bioinformatics. 2009;25:1754–60.

29. McKenna A, Hanna M, Banks E, Sivachenko A, Cibulskis K, Kernytsky A, et al. The Genome Analysis Toolkit: a MapReduce framework for analyzing next-generation DNA sequencing data. Genome Res. 2010;20:1297–303.

30. Robinson JT, Thorvaldsdóttir H, Winckler W, Guttman M, Lander ES, Getz G, et al. Integrative genomics viewer. Nat Biotechnol. 2011;29:24–6.

31. McLaren W, Gil L, Hunt SE, Riat HS, Ritchie GRS, Thormann A, et al. The Ensembl Variant Effect Predictor. Genome Biol. 2016;17:122.

32. Seabold S, Perktold J. Statsmodels: Econometric and Statistical Modeling with Python. Proceedings of the 9th Python in Science Conference. 2010;:92–6.

33. van Rossum G. Python library reference. 1995. https://ir.cwi.nl/pub/5009. Accessed 20 Jun 2020

