## Supplemental File 2 for "A nonsense mutation of bone morphogenetic protein-15 (BMP15) causes both infertility and increased litter size in pigs"

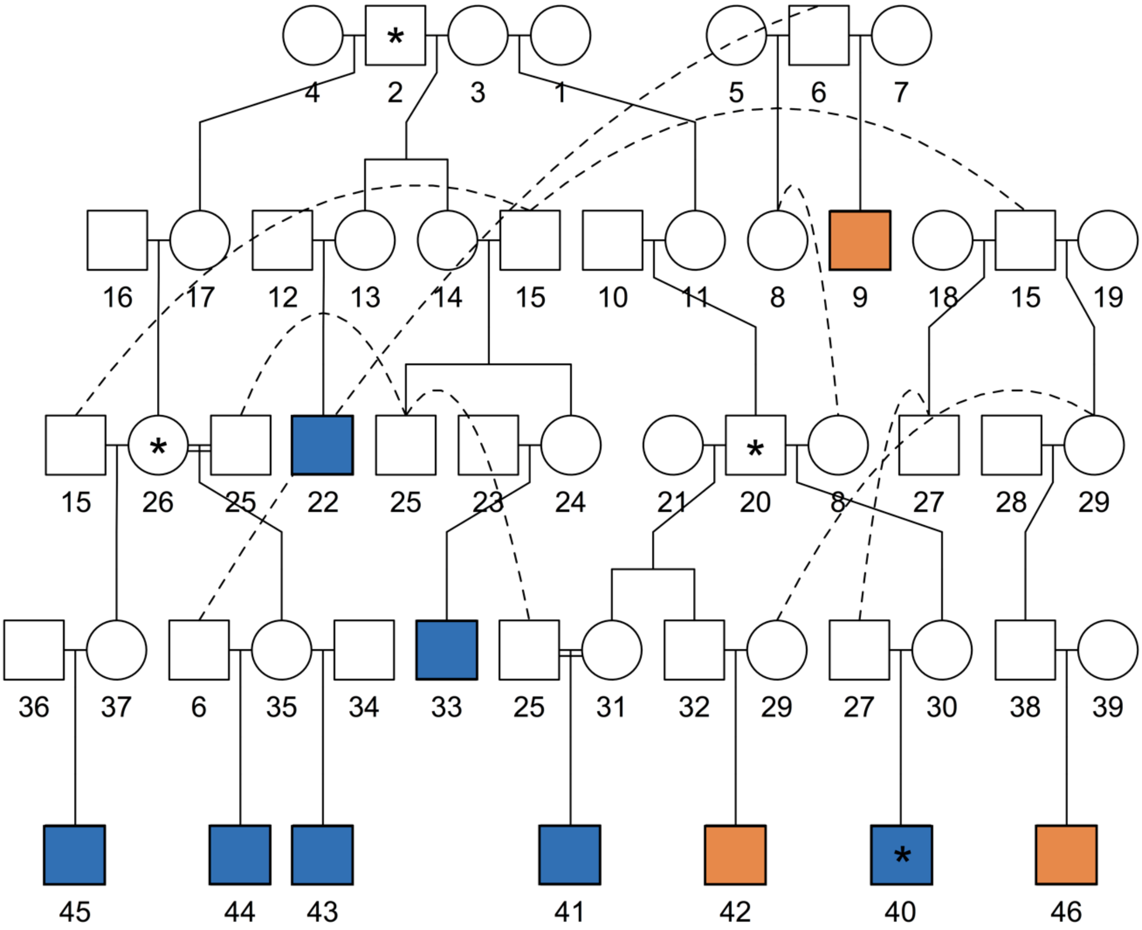


Pedigree listing sires of gilts with atypically small vulvae. Circles and rectangles represent females and males, respectively. Filled symbols denote sires of gilts that were classified as affected. The blue color denotes sires with the T-allele, leading to the nonsense mutation of BMP15 ( BMP15:p.R212X); the orange color symbolizes sires of gilts classified as affected but turning out to be CC. Symbols with asterisk denote animals with NGS-information.
