## Supplemental File 3 for "A nonsense mutation of bone morphogenetic protein-15 (BMP15) causes both infertility and increased litter size in pigs"

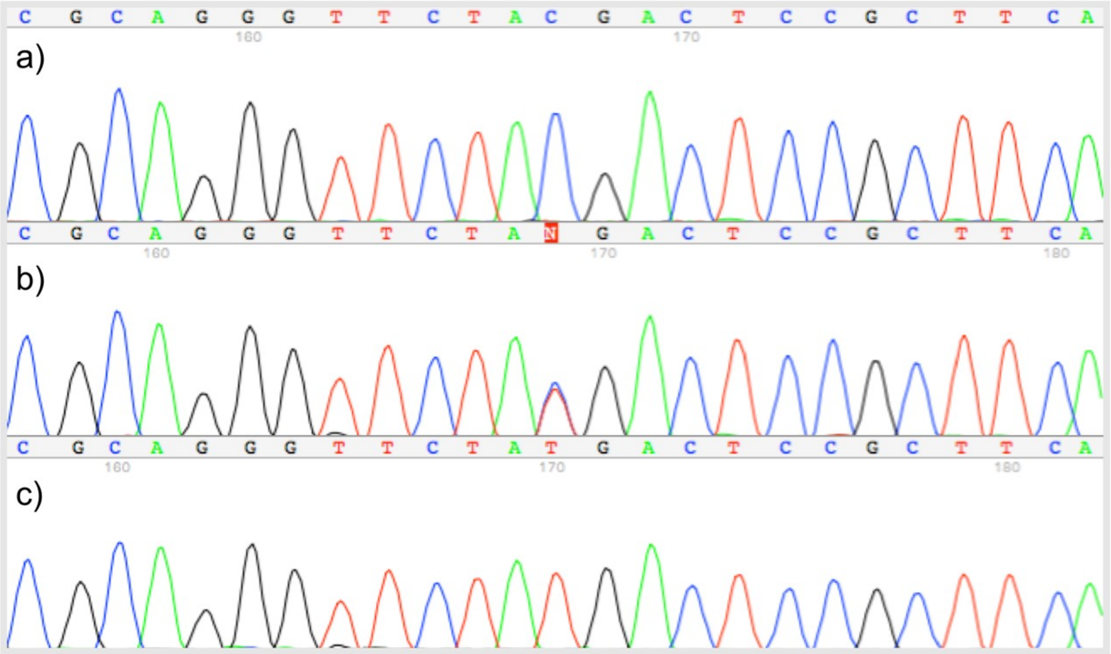


Chromatogram from Sanger-sequencing. a) homozygous for the wild type allele (C/C); b) heterozygous (C/T); c) homozygous for the mutant allele (T/T)
